# Human FcRn expression and Type I Interferon signaling control Echovirus 11 pathogenesis in mice

**DOI:** 10.1101/2020.09.24.312058

**Authors:** Alexandra I. Wells, Kalena A. Grimes, Kenneth Kim, Emilie Branche, Christopher J. Bakkenist, William H. DePas, Sujan Shresta, Carolyn B. Coyne

**Author notes:** Address correspondence: Carolyn Coyne, PhD, 9116 Rangos Research Center, UPMC Children’s Hospital of Pittsburgh, One Children’s Hospital Way, 4401 Penn Avenue, Pittsburgh, PA 15224.

## Abstract

Neonatal echovirus infections are characterized by severe hepatitis and neurological complications that can be fatal. Here, we show that expression of the human homologue of the neonatal Fc receptor (hFcRn), the primary receptor for echoviruses, and ablation of type I interferon (IFN) signaling are key host determinants involved in echovirus pathogenesis. We show that expression of hFcRn alone is insufficient to confer susceptibility to echovirus infections in mice. However, expression of hFcRn in mice deficient in type I interferon (IFN) signaling, hFcRn-IFNAR^−/−^, recapitulate the echovirus pathogenesis observed in humans. Luminex-based multianalyte profiling from E11 infected hFcRn-IFNAR^−/−^ mice revealed a robust systemic immune response to infection, including the induction of type I IFNs. Furthermore, similar to the severe hepatitis observed in humans, E11 infection in hFcRn-IFNAR^−/−^ mice caused profound liver damage. Our findings define the host factors involved in echovirus pathogenesis and establish *in vivo* models that recapitulate echovirus disease.

## Introduction

Echoviruses are small (∼30 nm) single-stranded RNA viruses that belong to the *Picornaviridae* family. Echoviruses consist of approximately 30 serotypes and are members of the Enterovirus genus, which are primarily transmitted through the fecal-oral route. Infants and neonates are often most severely impacted by echovirus infections, with the majority of enterovirus infections in infants below the age of two months caused by echoviruses^1,2^. Echovirus infections are particularly devastating in Neonatal Intensive Care Unit (NICU) outbreaks, where they account for 15-30% of nosocomial viral infections and can result in death of the neonate in as many as 25% of cases^3–6^. Echovirus 11 (E11) is one of the most common serotypes associated with outbreaks in NICUs across the world^7,8^. Despite the severe clinical outcomes associated with echovirus infections, the tissue tropism and pathogenesis of infection remain largely unknown due to the lack of established animal models to study E11 infection at secondary sites of infection, such as the liver and brain.

We and others previously identified the neonatal Fc receptor (FcRn) as a primary receptor for echoviruses^9,10^. Structural analysis has shown that the murine homologue of FcRn (mFcRn) does not support echovirus binding and entry^10^, which has also been shown experimentally in murine-derived primary cells and cell lines^9^. However, ectopic expression of human FcRn (hFcRn) renders murine-derived primary cells susceptible to echovirus infections^9^. FcRn is important for establishing passive immunity from mother to child through IgG transport across the placenta during human pregnancy or across the small intestine after birth in mice^11^. Additionally, FcRn is important for albumin homeostasis in liver hepatocytes and regulates the response to hepatic injury^12^. FcRn expression is maintained throughout life in the liver and many other tissue types in the body^13^. We have previously demonstrated in an oral infection model of suckling mice that E11 disseminates from the gastrointestinal (GI) tract into the blood and liver, and that this dissemination is dependent on the expression of human FcRn^9^. Although the virus disseminated to the liver, very little detectable virus was observed in this and other tissues, occluding further studies of pathogenesis at secondary sites of infection.

The development of mouse models that recapitulate the hallmarks of enterovirus disease in humans has historically been challenging. Enteroviruses typically do not infect mice as the murine homolog of their receptors are often not sufficient for binding and entry. Others have developed mouse models of select enteroviruses including poliovirus, coxsackievirus B (CVB), and enterovirus 71 (EV71)^14–16^. These models often use immunodeficient humanized transgenic mice, which express the human homolog of the receptor while lacking expression of the interferon α/β receptor (IFNAR)^15–19^. Despite established *in vivo* models for other enteroviruses, echoviruses have few established mouse models. A previous echovirus 1 mouse model was established using transgenic mice expressing human integrin very late antigen 2 (VLA-2), the receptor for E1^20^, which inoculated newborn mice intracerebrally, resulting in paralysis of the transgenic mice^21^. However, the host determinants involved in restricting echovirus infections *in vivo* remain largely unknown.

Here, we define the host determinants of echovirus infection and developed parallel adult and suckling mouse models of E11 infection. We show that immunocompetent animals that express hFcRn under the native human promotor (hFcRn^Tg32^) are largely resistant to E11 infection following intraperitoneal (IP) inoculation. In addition, immunodeficient mice lacking IFNAR expression (IFNAR^−/−^) alone are also refractory to infection. In contrast, hFcRn^Tg32^ animals that are also deficient in IFNAR expression (hFcRn^Tg32^-IFNAR^−/−^) are highly permissive to E11 infection and high levels of viral replication occur in the liver and pancreas, which reflects the tissue sites most commonly targeted in infected human neonates^22,23^. Luminex-based multianalyte profiling of whole blood revealed that hFcRn^Tg32^-IFNAR^−/−^ infected animals induced a robust systemic immune response to infection, including high levels of type I IFNs. Using RNASeq-based transcriptional profiling, we also show that the livers of hFcRn^Tg32^-IFNAR^−/−^ mice mount a pro-inflammatory and antiviral signaling cascade in response to infection. Finally, using hybridization chain reaction (HCR) with specific probes against the E11 genome, we show that hepatocytes are the main cell type infected in the liver. Our data thus define hFcRn and type I IFN signaling as key host determinant of E11 pathogenesis in the liver and suggest that these factors could be targeted therapeutically to control infection.

## Results

### Human FcRn and Type I IFN signaling are key host determinants of E11 infection

Given that the most severe outcomes of E11 infections in humans are in neonates, we first performed studies in suckling (7 day old) mice. We inoculated immunocompetent wild-type C57BL/6 (WT) and hFcRn^Tg32^ suckling mice with 10^4^ plaque forming units (PFU) of E11 by the IP route. Animals were sacrificed at 72 hours post inoculation (hpi) and tissues were collected for viral titration by plaque assay. Because an IP echovirus mouse model has not been established previously, we collected a diverse range of tissues (e.g. brain, liver, pancreas, small intestine) to determine the tissue tropism of E11 *in vivo*. WT and hFcRn^Tg32^ animals exhibited low to undetectable levels of infection in all of the tissues tested (**Figure 1A-F**). For example, only 2 of 12 WT animals and 2 of 13 hFcRn^Tg32^ animals had any detectable virus in liver and 0 of 12 WT mice and 1 of 13 hFcRn^Tg32^ mice had detectable virus in the brain, although in both cases, viral titers were very low **(Figure 1B, 1F)**. Because many enteroviruses are restricted by type I IFN signaling in small animal models and because we have previously shown that E11 is sensitive to recombinant IFN-β treatment^24^, we reasoned that type I IFNs might play a key role in restricting E11 infection *in vivo*. To test this, we infected suckling mice deficient in type I IFN signaling (IFNAR^−/−^) with 10^4^ PFU E11 by the IP route. However, we found that these animals were also largely resistant to E11 infection, with most animals having no detectable circulating virus in blood or replicating virus in tissues (4 of 12 animals had detectable virus in the blood and liver) **(Figure 1A-F)**. These data show that expression of hFcRn or ablation of type I IFN signaling alone is insufficient to confer susceptibility to E11 replication.

**Figure 1.**
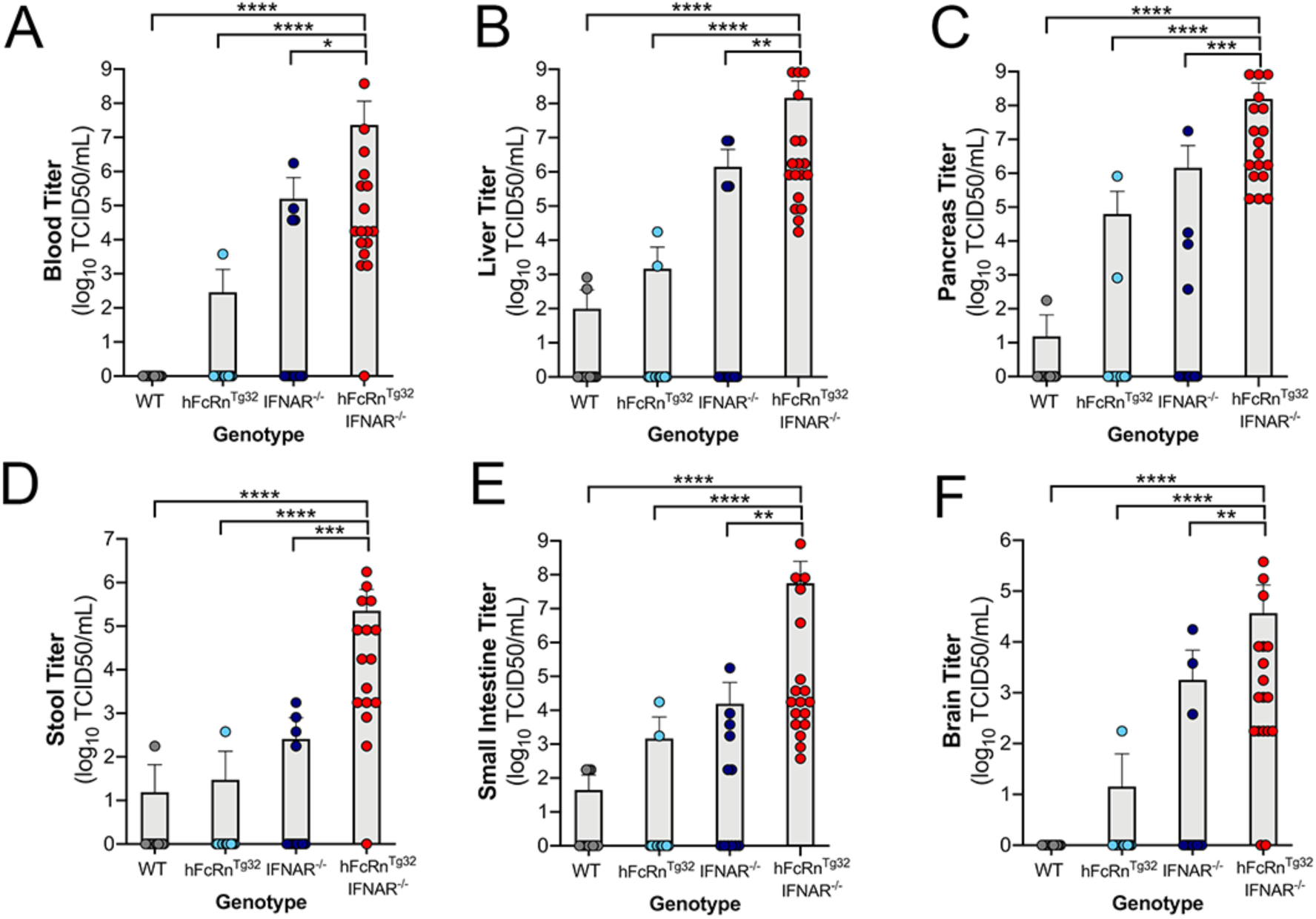
hFcRn^Tg32^-IFNAR^−/−^ suckling mice are permissive to E11 infection. C57Bl/6 (WT, gray), hFcRn^Tg32^ (light blue), IFNAR^−/−^ (dark blue), or hFcRn^Tg32^-IFNAR^−/−^ (red) suckling mice were IP inoculated with 10^4^ PFU of E11 and sacrificed 72 hours post inoculation. Viral titers (log_10_TCID50/mL) of suckling mice (WT – 12, hFcRn^Tg32^ – 13, IFNAR^−/−^ – 12, hFcRn^Tg32^-IFNAR^−/−^ – 18 animals) in the blood (A), liver (B), pancreas (C), stool (D), small intestine (E), and brain (F) are shown as mean ± standard deviation and individual animals (points). Data are shown with significance determined with a Kruskal-Wallis test with a Dunn’s test for multiple comparisons (*p<0.05, **p<0.005, ***p<0.0005, ****p<0.0001).

We next determined whether expression of hFcRn in the context of ablation of IFNAR-mediated signaling would be sufficient for E11 infection in mice. To do this, we generated hFcRn^Tg32^ mice that are deficient in IFNAR expression (hFcRn^Tg32^-IFNAR^−/−^). Similar to the studies described above, we inoculated suckling hFcRn^Tg32^-IFNAR^−/−^ mice with E11 by IP inoculation. In contrast to animals expressing hFcRn or lacking IFNAR expression alone, we found that hFcRn^Tg32^-IFNAR^−/−^ suckling mice were highly permissive to E11 infection, with high levels of infectious virus circulating in blood (17 of 18 animals, **Figure 1A)**. Similarly, hFcRn^Tg32^-IFNAR^−/−^ animals had significantly more detectable infectious virus in livers compared to other genotypes (18 of 18 with detectable virus in liver) **(Figure 1B)**. In addition to liver, we also observed high viral loads in the pancreas of hFcRn^Tg32^-IFNAR^−/−^ animals (18 of 18 with detectable virus, **Figure 1C)**. We also observed increased viral titers in the stool, small intestine, and brain, which all contained moderate to high levels of viral infection in hFcRn^Tg32^-IFNAR^−/−^ mice **(Figure 1D-F)**. These results show that hFcRn^Tg32^-IFNAR^−/−^ suckling mice are highly permissive to E11 inoculation.

We next determined whether hFcRn and IFN signaling played a role in echovirus pathogenesis in adult (6-week-old) mice. Similar to our findings in suckling mice, we found that WT, hFcRn^Tg32^, and IFNAR^−/−^ mice were largely resistant to E11 infection **(Figure 2A-F)**. In contrast to suckling mice, immunocompetent animals (WT and hFcRn^Tg32^) had no detectable circulating virus and a majority of IFNAR^−/−^ animals also completely resisted infection (2 of 16 with detectable virus in the blood) **(Figure 2A)**. In contrast, hFcRn^Tg32^-IFNAR^−/−^ animals had significant levels of viral replication in the blood (12 of 23 with detectable virus), liver (20 of 23 with detectable virus) and pancreas (13 of 23 with detectable virus), similar to what was observed in suckling pups **(Figure 2A-C)**. Additionally, these animals had low levels of detectable virus in the stool and small intestine suggesting this is not a main site of replication following IP inoculation **(Figure 2D & 2E)**. In contrast to suckling mice, adult hFcRn^Tg32^-IFNAR^−/−^ animals did not contain high levels of detectable virus in the brain (only 3 of 23 animals), suggesting age-related differences between adult and suckling mice **(Figure 2F)**. Taken together, these data show that both hFcRn and type I IFNs are key regulators of E11 infection of suckling mice and adult mice and that the liver is a key target site of replication *in vivo*.

**Figure 2.**
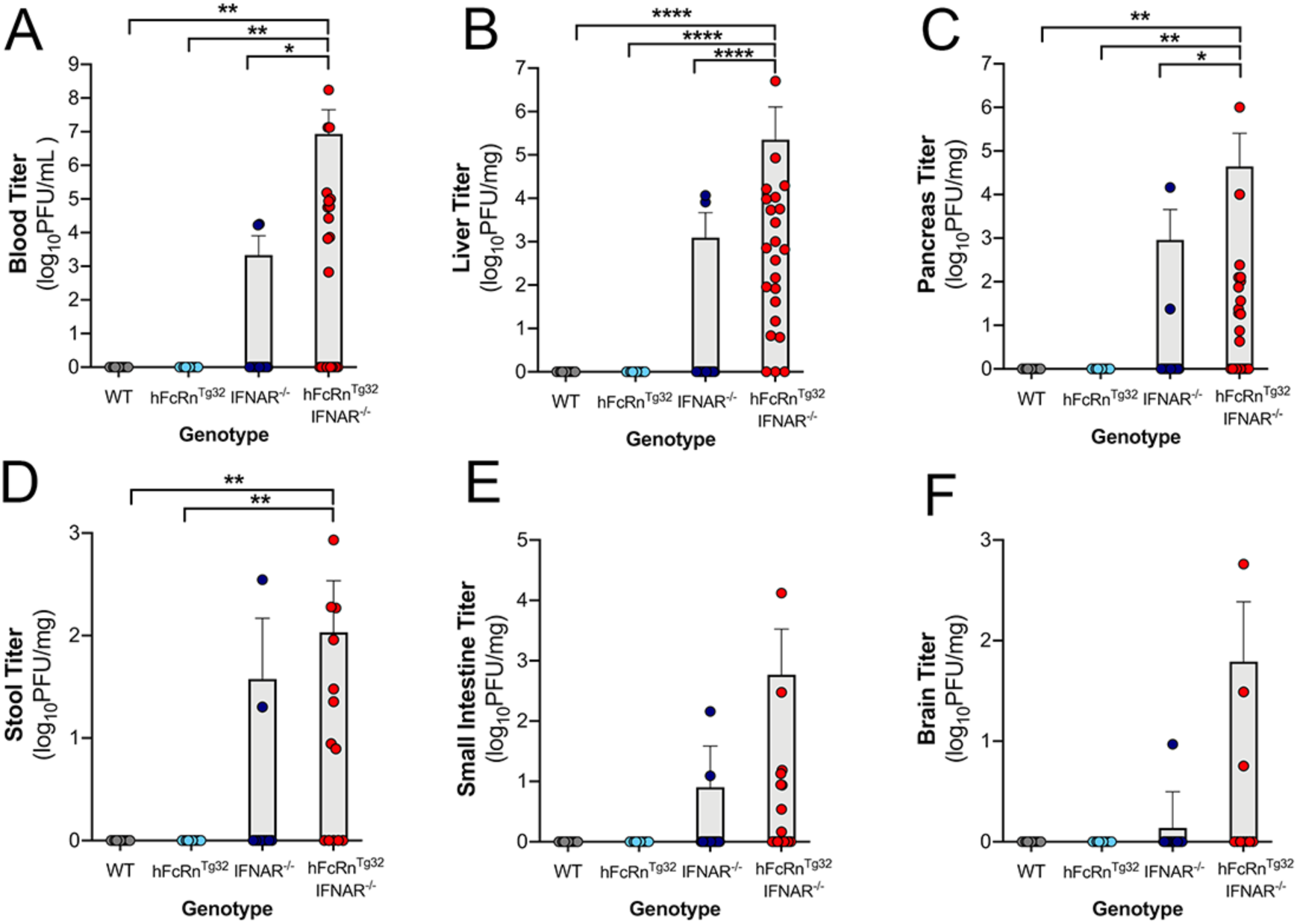
hFcRn^Tg32^-IFNAR^−/−^ adult mice are permissive to E11 infection. C57/BL6 (WT, gray), hFcRn^Tg32^ (light blue), IFNAR^−/−^ (dark blue), and hFcRn^Tg32^-IFNAR^−/−^ (red) animals were IP inoculated with 10^4^ PFU of E11 and sacrificed 72 hours post inoculation. **(A)** Viral titers in the blood (log_10_PFU/mL) of adult animals (WT – 11, hFcRn^Tg32^ – 10, IFNAR^−/−^ – 16, hFcRn^Tg32^-IFNAR^−/−^ – 23 animals). Viral titers in the liver (B), pancreas (C), stool (D), small intestine (E), and brain (F) (log_10_PFU/mg) from adult mice are shown as mean ± standard deviation bars and individual animals (points). Data are shown with significance determined with a Kruskal-Wallis test with a Dunn’s test for multiple comparisons (*p<0.05, **p<0.005, ***p<0.0005, ****p<0.0001).

### hFcRn^Tg32^-IFNAR^−/−^ animals induce a robust proinflammatory immune response to E11 infection

Due to the high levels of viremia in adult hFcRn^Tg32^-IFNAR^−/−^ mice, we next characterized the systemic immune response to E11 infection in these animals. To do this, we performed Luminex-based multiplex assays to assess the levels of 45 circulating cytokines and chemokines in the blood of adult animals infected with E11. Consistent with their low levels of infection, we observed no significant changes in the levels of circulating cytokines and chemokines in immunocompetent (WT, hFcRn^Tg32^) or immunodeficient (IFNAR^−/−^) mice (**Figure 3A**). In contrast, the blood of infected hFcRn^Tg32^-IFNAR^−/−^ animals contained high levels of various cytokines and chemokines in response to infection, with 19 cytokines/chemokines induced ≥ 2-fold compared to uninfected controls **(Figure 3A)**. The two most induced cytokines were members of the type I IFN family, IFN-α and IFN-β. On average, 7,802pg/mL of IFN-β was circulating in the blood of hFcRn^Tg32^-IFNAR^−/−^ animals, while WT, hFcRn^Tg32^, and IFNAR^−/−^ animals had little to no circulating IFN-β **(Figure 3B)**. Similarly, hFcRn^Tg32^-IFNAR^−/−^ animals had an average of 165pg/mL circulating IFN-α in blood while WT, hFcRn^Tg32^, and IFNAR^−/−^ animals had very low to undetectable levels **(Figure 3C)**. In addition to type I IFN induction, a number of chemokines, including monocyte chemoattractant protein 1 (MCP-1/CCL2), B cell attracting chemokine 1 (BCA-1/CXCL13), IP-10/CXCL10, and IL-12(p40) were present at very high levels in E11 infected hFcRn^Tg32^-IFNAR^−/−^ mice (**Figure 3D-G**). These data show adult hFcRn^Tg32^-IFNAR^−/−^ animals mount a potent immune response, including very high levels of type I IFNs, in response to E11 infection.

**Figure 3.**
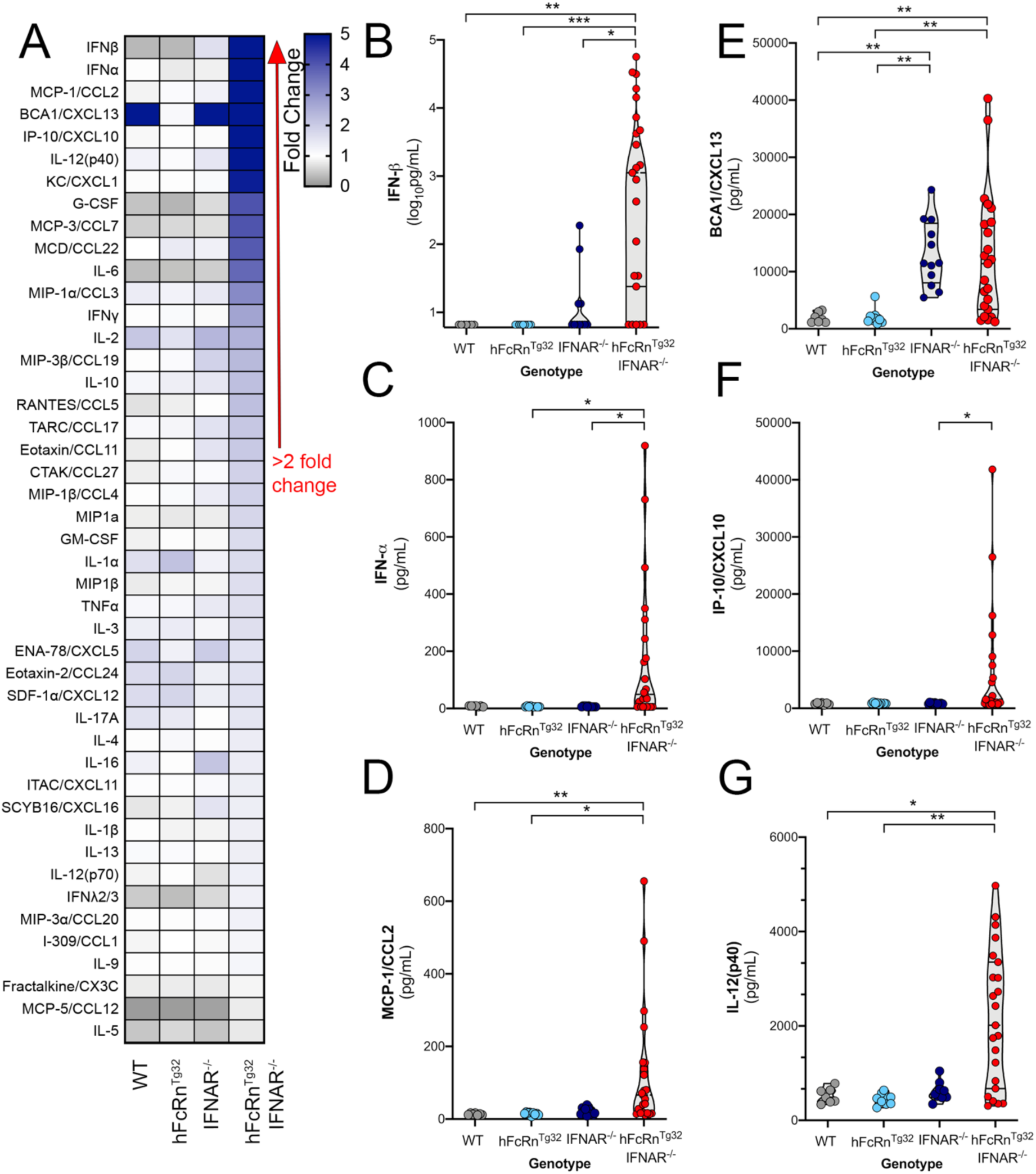
hFcRn^Tg32^-IFNAR^−/−^ animals induce a robust immune response to E11 infection. C57/BL6 (WT, gray), hFcRn^Tg32^ (light blue), IFNAR^−/−^ (dark blue), and hFcRn^Tg32^-IFNAR^−/−^ (red) animals were IP inoculated with 10^4^ PFU of E11 and sacrificed 72 hours post inoculation. Luminex-based multianalyte profiling of 45 cytokines was then performed from whole blood. **(A)** Heatmap demonstrating the induction (shown as fold-change from uninfected control) in E11-infected mice of the indicated genotype. Blue denotes significantly increased cytokines in comparison to untreated. Grey or white denote little to no changes (scale at top right). The red arrow demonstrates cytokines with greater than 2-fold upregulation observed in the average of separate experiments. Luminex assays were performed in duplicate. **(B-G)** IFN-β (B), IFN-α (C), MCL-1/CCL2 (D), BCA1/CXCL13 (E), IP-10/CXCL10, and IL12(p40) cytokine levels in the blood of E11 infected C57Bl/6 (WT, gray), hFcRn^Tg32^ (light blue), IFNAR^−/−^ (dark blue), and hFcRn^Tg32^-IFNAR^−/−^ (red) animals. Symbols represent individual mice. Significance was determined with a Kruskal-Wallis test with a Dunn’s test for multiple comparisons (*p<0.05, **p<0.005, ***p<0.0005).

### Infection and immune responses peak at 72h post-inoculation

Next, we determined the kinetics of the immune responses to E11 infection in hFcRn^Tg32^-IFNAR^−/−^ mice. To do this, we infected hFcRn^Tg32^-IFNAR^−/−^ animals with E11 and sacrificed at either 24, 48, or 72hpi and measured viral titers by plaque assays and immune induction by Luminex-based multiplex assays for 34 cytokines and chemokines. We found that there were measurable levels of virus present in key target tissues such as the blood, liver and pancreas by as early as 24hpi, with levels peaking at 72hpi **(Figure 4A-D)**. Consistent with these kinetics, we found that the levels of circulating cytokines increased at 24hpi and peaked at 72hpi as assessed by multianalyte Luminex-based profiling (**Figure 4E**). Strikingly, IFN-β was induced over ∼1,000pg/mL in animals infected for 24hrs and even higher in animals after 48hpi and 72hpi **(Figure 4F)**. In addition, IFN-α and IFN-λ2/3 were increased at 72hpi compared to control and 24hpi **(Figure 4G & 4H)**. In contrast to IFNs, other cytokines and chemokines including IP-10/CXCL10, MCP-1/CCL2, and KC/CXCL1 were induced at highest levels at 48hpi, with levels decreasing by 72hpi (**Figure 4I-K**). These data suggest that animals induce an immune response to infection very early following the initiation of viral replication.

**Figure 4.**
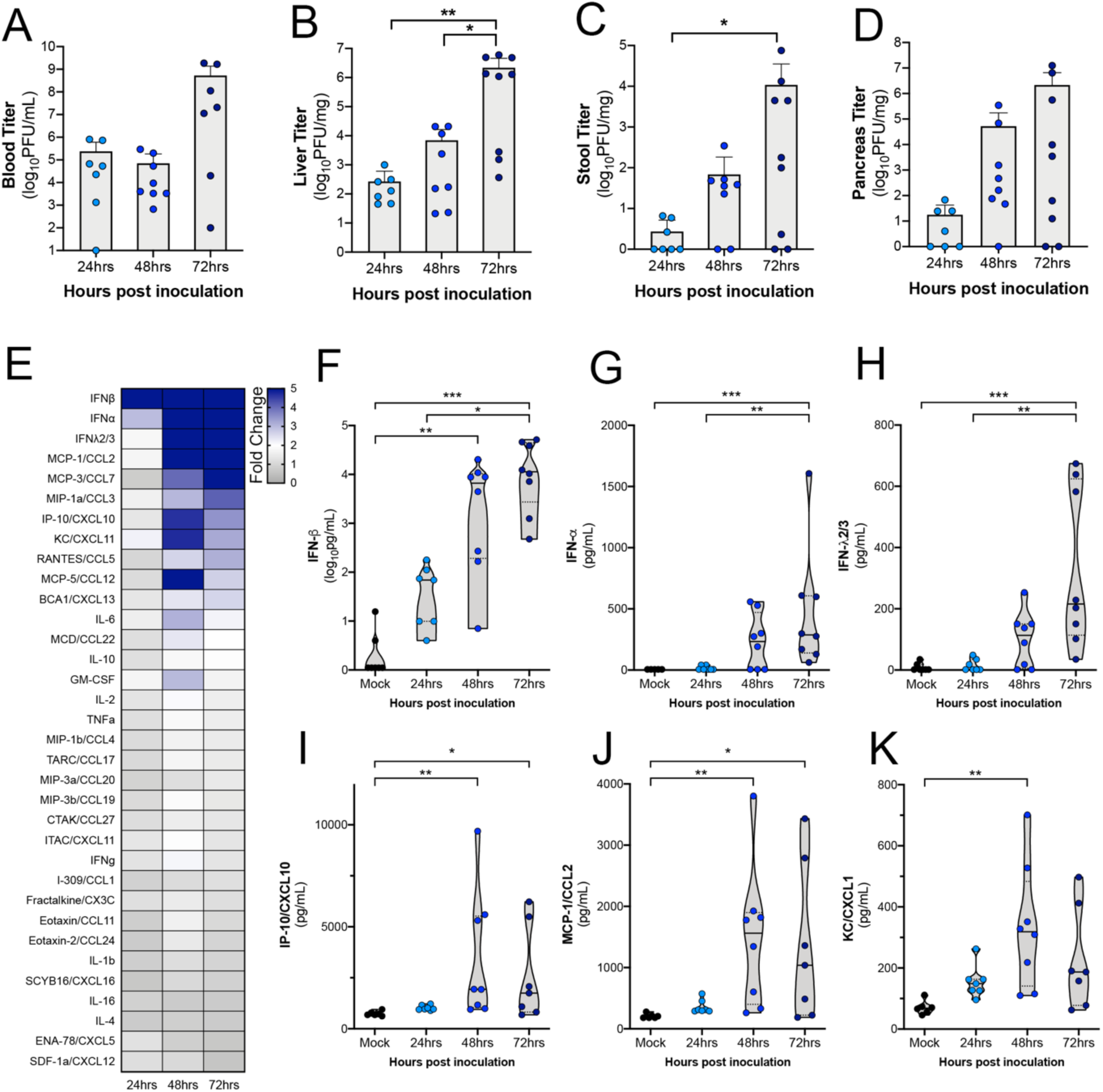
Cytokine levels increase with viremia in hFcRn^Tg32^-IFNAR^−/−^ animals. hFcRn^Tg32^-IFNAR^−/−^ animals IP inoculated with 10^4^ PFU of E11 were sacrificed at 24 (light blue) 48 (blue), or 72 (navy) hours post inoculation. **(A)** Viral titers in the blood (log_10_PFU/mL) of adult animals (24hpi – 7, 48hpi – 8, 72hpi – 9 animals) are shown as mean ± standard deviation bars) and individual animals (points). **(B-D)** Viral titers in the liver (B), stool (C), and pancreas (D), (log_10_PFU/mg) from adult mice are shown as mean ± standard deviation bars and individual animals (points). **(E)** Heat map demonstrating the level of protein induction by Luminex-based assays shown as the fold change of from the average pg/mL of the uninfected animals to each individual animal concentration per protein then averaged within each timepoint. Proteins are sorted from largest fold change (blue) from uninfected to smallest fold change (gray) in 72hpi animals. **(F-K)** IFN-β (F), IFN-α (G), IFNλ2/3 (H), IP-10/CXCL10 (I), MCP-1/CCL2 (J), and KC/CXCL1 (K) protein levels expressed in the blood of each animal shown by timepoint. Data are shown with significance determined with a Kruskal-Wallis test with a Dunn’s test for multiple comparisons (*p<0.05, **p<0.005, ***p<0.0005, ****p<0.0001).

### E11 infection induces damage and cell death in the livers of hFcRn^Tg32^-IFNAR^−/−^ animals

Echovirus infections in neonates commonly induces liver failure, which can be fatal^23^. In addition, our data suggested that the highest levels of E11 replication in hFcRn^Tg32^-IFNAR^−/−^ mice was in the liver. Thus, we focused on the impact of E11 infection on the liver as a contributor to disease. Blinded pathology scoring of H&E stained sections of infected livers revealed no histopathologic changes in immunocompetent animals or in IFNAR^−/−^ adult or suckling mice infected with E11 (**Figure 5A-B, Supplemental Figure 1**). In contrast, there was moderate to severe liver damage induced by E11 infection of adult hFcRn^Tg32^-IFNAR^−/−^ animals, including punctate hepatocytolysis and necrosis at 72hpi **(Figure 5A, Supplemental Figure 1)**. Other histopathological changes included increased immune cell infiltration, which was also observed in infected hFcRn^Tg32^-IFNAR^−/−^ suckling mice (**Figure 5B**, black arrows). In addition to histopathology, we assessed the impact of infection on cell viability using an antibody specific for the cleaved (activated) version of caspase-3. Whereas E11 infection of immunocompetent and IFNAR^−/−^ animals exhibited no cleaved caspase-3 staining as assessed by immunohistochemistry, E11-infected hFcRn^Tg32^-IFNAR^−/−^ adults and suckling mice exhibited pronounced positive cleaved caspase-3 staining **(Figure 5C, 5D)**. These data indicate that the livers of hFcRn^Tg32^-IFNAR^−/−^ animals undergo apoptosis and cell death following E11 infection.

**Figure 5.**
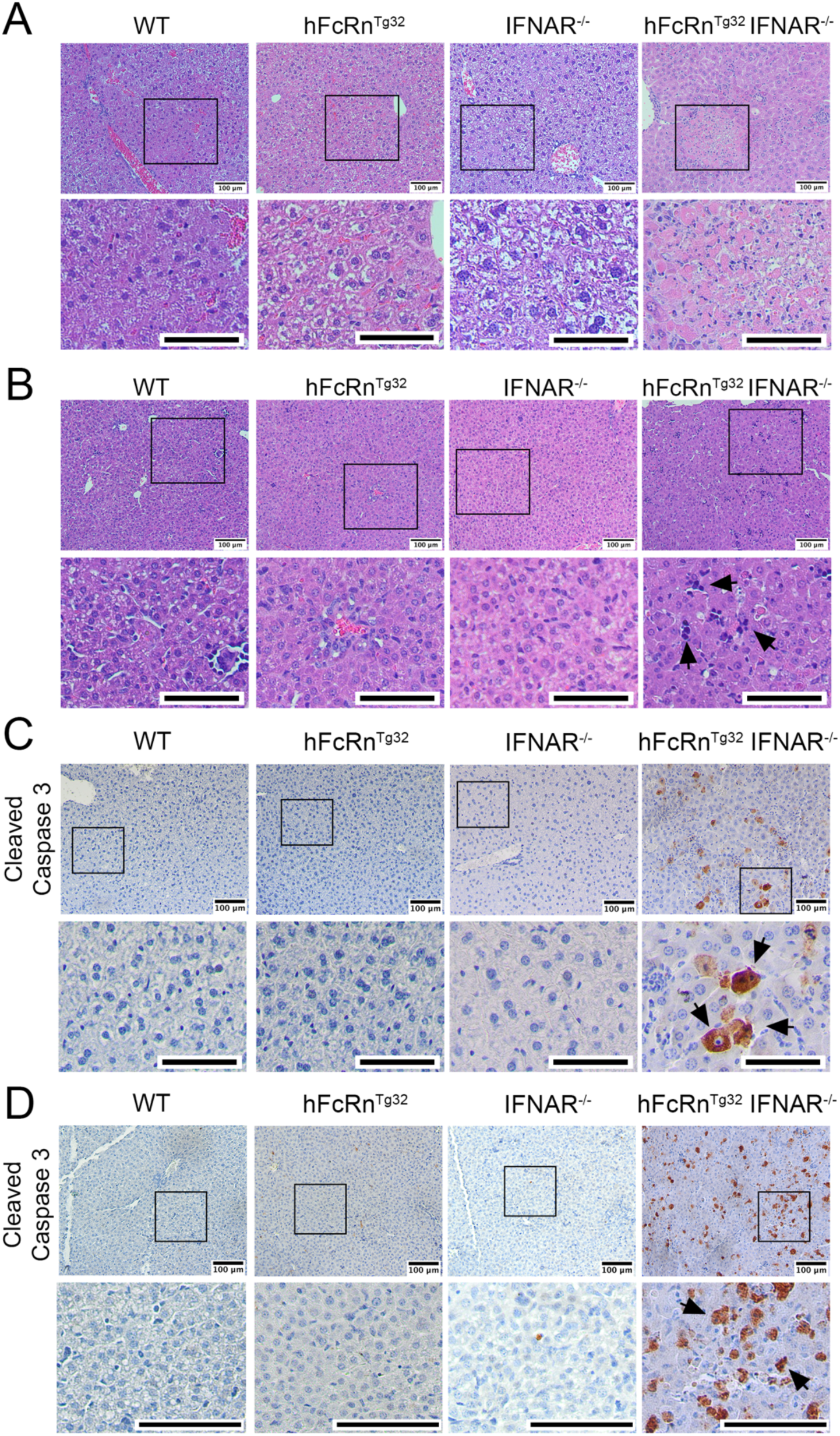
E11 infection induces histopathologic changes and cell death. C57Bl/6 (WT), hFcRn^Tg32^, IFNAR^−/−^, and hFcRn^Tg32^-IFNAR^−/−^ adult **(A & C)** or suckling mice **(B & D)** were IP inoculated with 10^4^ E11 and sacrificed 72 hours post inoculation. **(A & B)** H&E staining of the livers in adult (A) or suckling (B) mice. **(C & D)** Immunohistochemistry using an antibody recognizing the cleaved form of caspase 3 from the livers of a representative animal of each genotype as indicated. Adult (C) and suckling mice (D). Black arrows denote positive staining. Scale bars (100μm) are shown at bottom right.

### E11 infection of hFcRn^Tg32^-IFNAR^−/−^ mice induces a robust local proinflammatory immune response in the liver

Because we found that the livers of hFcRn^Tg32^-IFNAR^−/−^ mice infected with E11 exhibited histopathologic changes and underwent cell death, we profiled other liver changes by RNAseq transcriptional profiling. Consistent with our Luminex-based profiling studies of circulating cytokines, we found that the livers of hFcRn^Tg32^-IFNAR^−/−^ animals infected with E11 robustly induced expression of the transcripts for type I IFNs, with less robust induction of type III IFNs (**Figure 6A**). Levels of vRNA in infected animals mirrored our findings on infectious viral titers, with high levels in hFcRn^Tg32^-IFNAR^−/−^ mice (**Figure 6B)**. In addition to these changes, hFcRn^Tg32^-IFNAR^−/−^ infected animals also induced the expression of other pro-inflammatory and immunomodulatory factors, including chemokines (e.g. Ccl2, Cxcl1, Cxcl9), transcription factors (e.g. Stat1, Stat3, Socs1), and interferon stimulated genes (e.g. Isg15, Ifit1) (**Figure 6C, D**).

**Figure 6.**
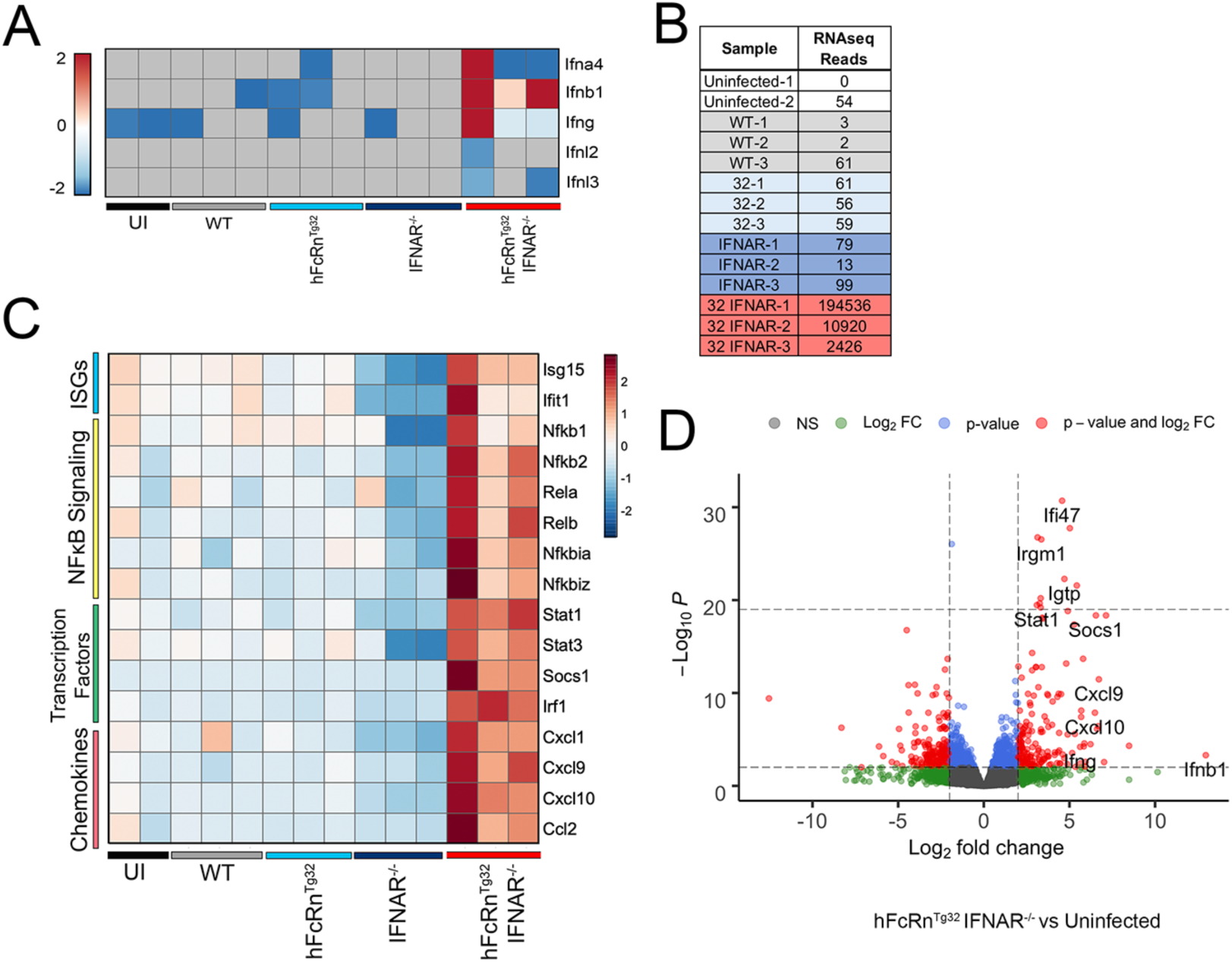
Transcriptional profiling from the livers of E11 infected hFcRn^Tg32^-IFNAR^−/−^ animals reveals induction of a proinflammatory immune response to infection. RNAseq-based transcriptional profiling from RNA isolated from the livers of E11 infected C57Bl/6 (WT), hFcRn^Tg32^, IFNAR^−/−^ or hFcRn^Tg32^-IFNAR^−/−^ animals (3 animals each), or uninfected controls (2 animals) was performed. **(A)** Heatmap of log_2_RPKM values for type I (Ifna4, Ifnb1), II (Ifng), and III (Ifnl2, Ifnl3) IFNs in the livers of the indicated genotypes 72hpi. Scale shown at left. **(B)** RPKM values mapped to the E11 genomic sequence in each genotype. Individual animals are shown. **(C)** Heatmap based on log_2_RPKM values of select proinflammatory cytokines in the livers of following E11 infection of the indicated genotypes, or uninfected controls. Scale is shown at right. In (A) and (C), red indicates higher expression and blue indicates lower expression. Grey denotes no reads detected. **(D)** Volcano plot of differentially regulated genes in hFcRn^Tg32^-IFNAR^−/−^ adult animals compared to uninfected animals. Red indicates genes with a statistically significant upregulation or downregulation of > or < log_2_ fold-change of 2 and p<0.05.

### E11 specifically infects hepatocytes in hFcRn^Tg32^-IFNAR^−/−^ mice

Finally, we defined the cellular tropism of E11 within the liver. Using immunohistochemistry for the viral VP1 capsid protein, we found that E11 localized primarily in what appeared to be hepatocytes **(Figure 7A)**. No positive staining for VP1 was observed in any other three mouse strains **(Figure 7A)**. hFcRn^Tg32^-IFNAR^−/−^ suckling mice also displayed positive VP1 staining in the liver **(Supplemental Figure 2)**. Although VP1 staining suggested that E11 replication occurred primarily in hepatocytes, we developed a more sensitive approach to define the cellular tropism of E11 using hybridization chain reaction (HCRv3.0). HCR allows for multiplexed quantitative RNA fluorescence *in situ* hybridization (RNA-FISH) and the signal amplification inherent to the technique vastly enhances the dynamic range and sensitivity of conventional FISH-based approaches^25–27^. To do this, we designed probes specific for the E11 genome and performed HCR on liver sections from hFcRn^Tg32^-IFNAR^−/−^ mice infected with E11 (schematic, **Figure 7B**). To define the localization of E11 specifically to hepatocytes, we also developed probes to albumin, a specific marker of hepatocytes. Using HCR, we observed the presence of E11 vRNA in the livers of infected mice by 24hpi, with the numbers of positive cells increasing by 48-72hpi (**Figure 7C**). Interestingly, E11 vRNA positive cells exclusively colocalized with albumin, identifying hepatocytes as the main cellular target of infection in the liver. To confirm this, we quantified three fields at each time point and quantified colocalization between vRNA and albumin signals, which revealed a strong colocalization (Pearson’s coefficient 24hpi – 0.73, 48hrs – 0.85, 72hpi – 0.84). Together, these data show that E11 replicates in liver hepatocytes in hFcRn^Tg32^-IFNAR^−/−^ animals.

**Figure 7.**
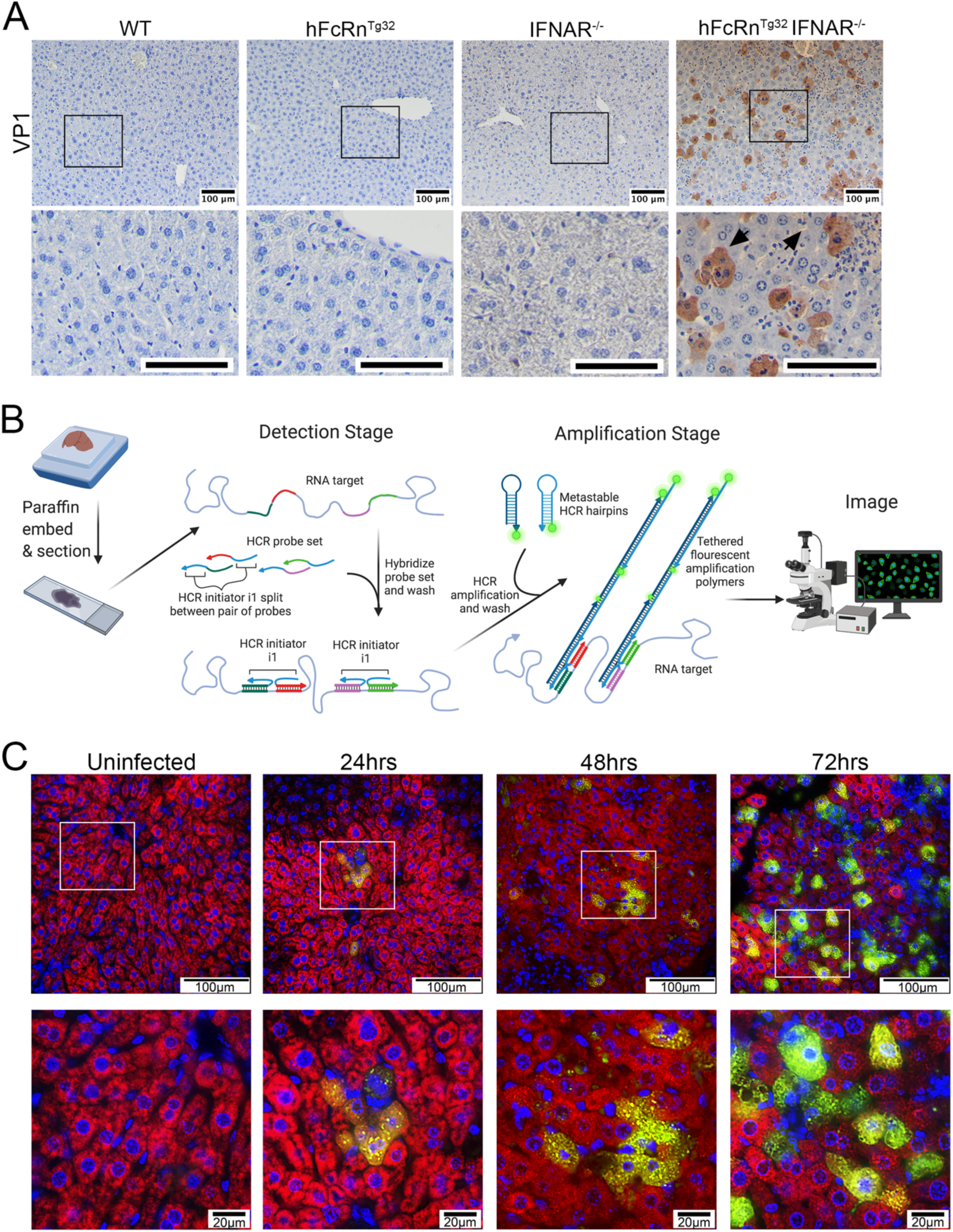
Hepatocytes are the primary site of E11 replication in the liver. **(A) C57Bl/6 (**WT), hFcRn^Tg32^, IFNAR^−/−^, or hFcRn^Tg32^ IFNAR^−/−^ adult animals were IP inoculated with 10^4^ PFU of E11 646 and sacrificed 72 hours post inoculation. Immunohistochemistry for E11 using an antibody 647 recognizing the VP1 capsid protein from the liver of a representative adult animal of each 648 genotype is shown. Black arrows denote positive staining. **(B)** Schematic of the hybridization 649 chain reaction (HCR) protocol used adapted from the Molecular Instruments HCR v3.0 protocol 650 and created with BioRender.com. **(C)** HCR of hFcRn^Tg32^-IFNAR^−/−^ adult animals at the indicated 651 dpi using probes against the E11 genome (green) and albumin (red). White boxes denote areas 652 zoomed at bottom. Scale bars shown at bottom right (100μm at top and 20μm at bottom). Three 653 unique fields were captured and colocalization between vRNA and albumin quantified, as 654 indicated in the text.

## Discussion

Here, we show that human FcRn and type I IFN signaling are key host determinants that control E11 infection in the liver, a tissue site commonly associated with human disease. Through Luminex-based multianalyte and RNASeq-based transcriptional profiling, we also show that animals expressing hFcRn and ablated in type I IFN signaling initiate a systemic immune response to infection. Furthermore, we show that E11 replication in the liver induces histopathological changes and apoptotic cell death in hepatocytes. Our findings thus define proviral (hFcRn) and antiviral (type I IFN) host factors that control echovirus infections specifically in the liver. In addition, our studies provide a novel animal model that can be used to test anti-echovirus therapeutics.

Although FcRn has been identified as a pan-echovirus receptor^9,10^, its role in mediating echovirus pathogenesis has remained unclear. Previous work has shown that FcRn is expressed in many different cell types in the body, including the small intestine^28,29^ and in liver hepatocytes^30,31^. Despite what its name implies, FcRn is expressed on many cells throughout life, often at very high levels. Our results shown here define the organs targeted by E11 in an *in vivo* model, with high levels of replication in various tissues, such as the liver and pancreas. Our parallel adult and suckling pup models allowed us to compare age-related differences that might impact sensitivity or responses to echovirus infections. Of note, the animals used in our studies express hFcRn under the control of the endogenous promoter, which might mimic age-related changes in expression observed in humans. Interestingly, although we detected high levels of echovirus replication in similar tissues between adults and suckling pups, there were age-related differences in viral infection in the brains of these mice. Whereas 16 of 18 of infected hFcRn^Tg32^-IFNAR^−/−^ suckling mice exhibited replication in the brain, only 3 of 23 adult animals did. Although this could be attributed to differences on the relative ratio of weight to viral inoculum, circulating viral titers in the blood were similar between suckling pups and adult mice. Given that echovirus infections are commonly associated with aseptic meningitis in neonates, these findings suggest that expression levels of hFcRn and type I IFN signaling could be key determinants of age-related susceptibility in key sites targeted in humans, such as the liver and brain.

The liver is a primary site of echovirus-associated disease, with hepatitis and acute liver failure commonly observed in infected infants and children and the majority of echovirus-associated death in neonates occurs due to overwhelming liver failure^32^. Our *in vivo* findings suggest that FcRn expression is required for this infection only when host type I IFN signaling is ablated. In addition to IFNs, we observed induction of a number of other immunomodulatory factors in infected animals. The role of cytokines in echovirus pathogenesis in humans is not known. However, immunodeficient individuals, including adults, are more susceptible to echovirus infections, which often induces hepatitis^33–36^. In addition, analysis of mutations in the E11 genome induced by selective pressure in an immunodeficient individual who developed chronic infection revealed strikingly high sequence conservation in the 3C virally-encoded protease which often attenuates host cell innate immune signaling^36^. Our studies suggest that type I IFNs are the primary drivers of resistance to echovirus infections in the liver, which is supported by our RNASeq studies, in which low levels of the transcripts for type III IFNs were upregulated by infection. These findings are similar to those for the related enterovirus coxsackievirus B3 (CVB3), whose infection in the liver is also regulated primarily by type I IFN signaling^19^. Collectively, our studies show that expression of hFcRn in the setting of diminished type I IFN signaling is the primary driver of E11 infection in the liver.

Despite the clear hepatic tropism of echoviruses, little is known regarding the cell type(s) targeted by echoviruses in the liver or how these cells respond to infection. Moreover, the role of FcRn in mediating this tropism is unknown. The liver is composed of diverse cell types. In addition to hepatocytes, which comprise ∼80% of total liver cells, tissue resident Kupffer Cells represent ∼35% and liver sinusoidal endothelial cells comprise ∼50% of non-parenchymal cells. FcRn is thought to be expressed in all of these cell types^37^. Our studies thus define the tropism of echoviruses specifically to hepatocytes and show that FcRn expression is a key determinant of this tropism. In addition, our studies suggest that echovirus infection of hepatocytes induces pronounced hepatic damage, characterized by apoptotic cell death and tissue damage. These findings are consistent with what is observed in autopsy tissue isolated from echovirus infected neonates, which also indicates extensive infection-induced hepatocyte damage^23,35,38,39^.

Consistent with high levels of infection in the liver, hFcRn^Tg32^-IFNAR^−/−^ infected animals also exhibited infectious virus present in the stool. Given that echoviruses are transmitted by the fecal-oral route, defining how viral particles are shed and subsequently transmitted is important for understanding pathogenesis and spread. Because infected animals did not have high titers in the small intestine (∼10^2^ PFU/mg on average), our data indicate that shed virus does not result from direct intestinal infection, which is expected given the route of inoculation. The most likely scenario is via the gut-liver axis. Many studies have shown that the bacteria and bacterial products can reach the liver through the portal vein and liver secretory products, such as bile acids, IgA, and antimicrobial molecules, can leave the liver into the intestines through the biliary tract^40,41^. It is thus likely that infectious virus exits the liver through the biliary tract into the intestine where it exits the body in the stool, explaining the high stool titers with little to no infectious virus in the intestine itself.

There are currently no effective antiviral therapeutics to combat echovirus infections. Our work thus establishes *in vivo* models that full recapitulate echovirus infection in human neonates and could thus be used to develop and test antivirals. In addition, our studies define key roles for FcRn and type I IFN signaling in mediating echovirus pathogenesis and suggest these factors could be targeted to ameliorate or prevent infections. Collectively, this work defines fundamental aspects of echovirus biology that enhance our understanding of how infection, tissue targeting, and disease occurs.

## Materials and Methods

### Cell lines and viruses

HeLa cells (clone 7B) were provided by Jeffrey Bergelson, Children’s Hospital of Philadelphia, Philadelphia, PA, and cultured in MEM supplemented with 5% FBS, non-essential amino acids, and penicillin/streptomycin. Experiments were performed with echovirus 11 Gregory (E11), which was obtained from the ATCC. Virus was propagated in HeLa cells and purified by ultracentrifugation over a 30% sucrose cushion, as described previously^42^.

### Animals

All animal experiments were approved by the University of Pittsburgh Animal Care and Use Committee and all methods were performed in accordance with the relevant guidelines and regulations. C57BL/6J (WT, cat. no. 000664), B6.Cg-*Fcgr*^*t*tm1Dcr^Tg(FCGRT)32Dcr/DcrJ (hFcRn^Tg32^, cat. no. 014565), B6(Cg)-Ifnar1^tm1.2Ees^/J (IFNAR^−/−^, cat. no. 028288) were purchased from The Jackson Laboratory. hFcRn^Tg32^-IFNAR^−/−^ mice were generated by crossing B6.129S2-Ifnar1^tm1Agt^/Mmjax (cat no. 32045-JAX) with B6.Cg-Fcgrt^tm1Dcr^ Tg(FCGRT)32Dcr/DcrJ (cat no. 014565). Breeders were established that were deficient in mouse FcRn and IFNAR and were homozygous for the hFcRn transgene. All animals used in this study were genotyped by Transnetyx.

### Adult animal infections

6-7-week-old mice were inoculated by the intraperitoneal route with 10^4^ PFU of E11. Intraperitoneal inoculation was performed using a 1mL disposable syringe and a 25-gauge needle in 100μL of 1X PBS. Mice were euthanized at 3 days post inoculation, or at times specified in the figure legends, and organs harvested into 1mL of DMEM (viral titration) or RNA lysis buffer (RNA isolation) and stored at −80°C. Tissue samples for viral titration were thawed and homogenized with a TissueLyser LT (Qiagen) for 8 minutes, followed by brief centrifugation for 5 minutes at 5000 x g. Viral titers in organ homogenates were determined by plaque assay in HeLa cells overlayed with a 1:1 mixture of 1% agarose and 2x MEM (4% FBS, 2% pen/strep, 2% NEAA). Plaques were enumerated 40hpi following crystal violet staining.

### Suckling pup infections

7-day-old mice were inoculated by the intraperitoneal route with 10^4^ PFU of E11. Two separate litters were inoculated for each condition. Intraperitoneal inoculation was performed using a 1mL disposable syringe and a 27-gauge needle in 50μL of 1X PBS. Mice were euthanized at 3 days post inoculation and organs harvested into 0.5mL of DMEM (viral titration) or RNA lysis buffer (RNA isolation) and stored at −80°C. Tissue samples for viral titration were thawed and homogenized with a TissueLyser LT (Qiagen) for 5 minutes, followed by brief centrifugation for 5 minutes at 8000 x g. Viral titers in organ homogenates were determined by TCID50 in HeLa cells and enumerated following crystal violet staining.

### Immunohistochemistry

Tissues were fixed in 10% buffered formalin for 24hrs and then transferred to 70% ethanol. Tissues were embedded in paraffin and sectioned. Slides were stained with a monoclonal VP1 antibody, as described previously^9^, or cleaved caspase 3. Tissue sections were deparaffinized with xylene and rehydrated with decreasing concentrations of ethanol (100%, 95%, 80%), then washed with ddH_2_0. Antigen unmasking was performed with slides submerged in 10 mM citrate buffer (pH 6.0) and heated in a steamer for 20 minutes at ∼90°C. Slides were cooled to room temperature and slides were immunostained with cleaved caspase 3 using Vectastain Elite ABC HRP (Vector Biolabs, PK-6100), according to the manufacturer’s instructions. Slides were incubated in 6% H_2_O_2_ in methanol for 30 min then washed 3 times for 5 minutes in H_2_O. Avidin block (Vector, SP-2001) was applied for 15 minutes and washed twice in H_2_O followed by biotin block (Abcam, ab156024) for 15 minutes and washed twice in H_2_O. Finally, serum-free protein block was applied for 10 minutes and cleaved caspase 3 antibody was diluted 1:100 in TBS-T (Tris-buffered saline, 0.1% Tween 20) and slides incubated overnight in a humidified chamber at 4C. Next, slides were washed three times for 5 min in PBST and exposed to the goat anti-rabbit biotinylated secondary antibody (Vector, BA-1000) for 30 min. Slides were rinsed in PBST three times for 5 min and the Vectastain Elite ABC HRP kit was applied for 30 min. Slides were rinsed in PBST for three times for 5 min and diaminobenzidine substrate for 5 mins; which was terminated with water incubation. Slides were counterstained with hematoxylin for 1 min, thoroughly rinsed with H_2_O, and incubated in 0.1% sodium bicarbonate in H2O for 5 mins. Slides were then dehydrated with increasing concentrations of ethanol, cleared with xylene and mounted with Cytoseal 60 (Thermo Scientific, 83104). Images were captured on an IX83 inverted microscope (Olympus) using a UC90 color CCD camera (Olympus).

### Antibodies

The following antibodies were used-anti-VP1 (NCL-ENTERO, clone 5-D8/1, Leica Biosystems) and cleaved caspase 3 (Asp175) (9661, Cell Signaling).

### HCR and Imaging

HCR was performed following the Molecular Instruments HCR v3.0 protocol for FFPE human tissue sections^25,27^. Briefly, tissue sections were deparaffinized with xylene and rehydrated with decreasing concentrations of ethanol (100%, 95%, 80%). Antigen unmasking was performed with slides submerged in 10 mM citrate buffer (pH 6.0) and heated in a steamer for 20 minutes at ∼90°C. Slides were cooled to room temperature. Sections were treated with 10 µg/mL Proteinase K for 10 min at 37°C and washed with RNase free water. Samples were incubated for 10 minutes at 37°C in hybridization buffer. Sections were incubated overnight in a humidified chamber at 37°C with 0.4 pmol of initiator probes in hybridization buffer (Table 1 echovirus probes, Table 2 albumin probes). The next day, slides were washed in probe wash buffer and 5x SSCT for 4x 15 min, according to the manufacturer’s instructions. Samples were incubated in a humidified chamber at 37°C for 30 minutes in amplification buffer. Fluorescent hair pins were heated to 95°C for 90 seconds and snap cooled at room temperature for 30 min. Hairpins and amplification buffer were added to the sample and incubated overnight at room temperature. Hairpins were washed off with 5x SSCT for 5 minutes, 15 minutes, 15 minutes, and 5 minutes. Slides were mounted in vectashield with DAPI. Slides were imaged an IX83 inverted microscope (Olympus) with ORCA-FLASH 4.0 camera. Olympus CellSens advanced imaging software with the deconvolution package, constrained iterative, was used.

**Table 1.**
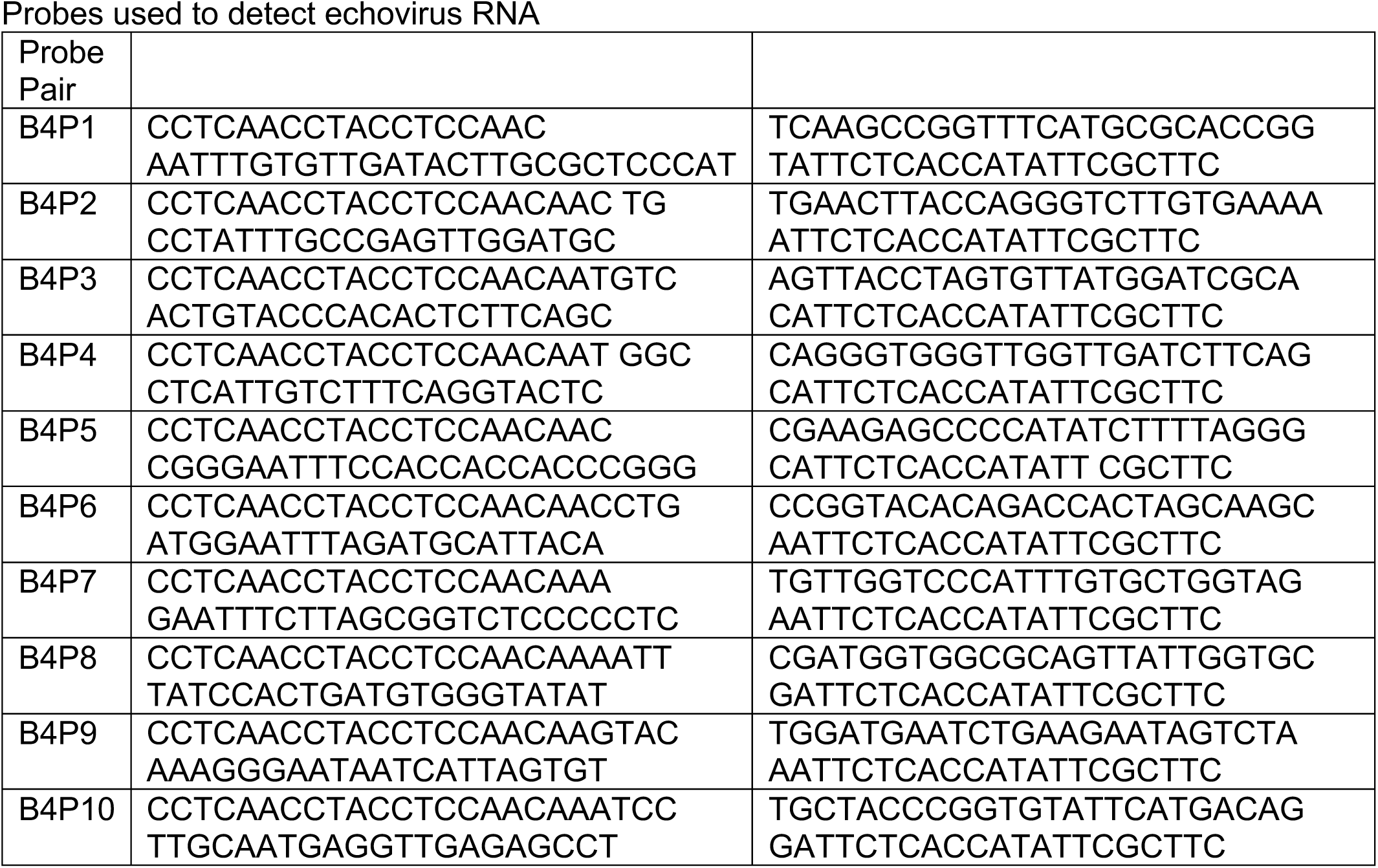

**Table 2.**
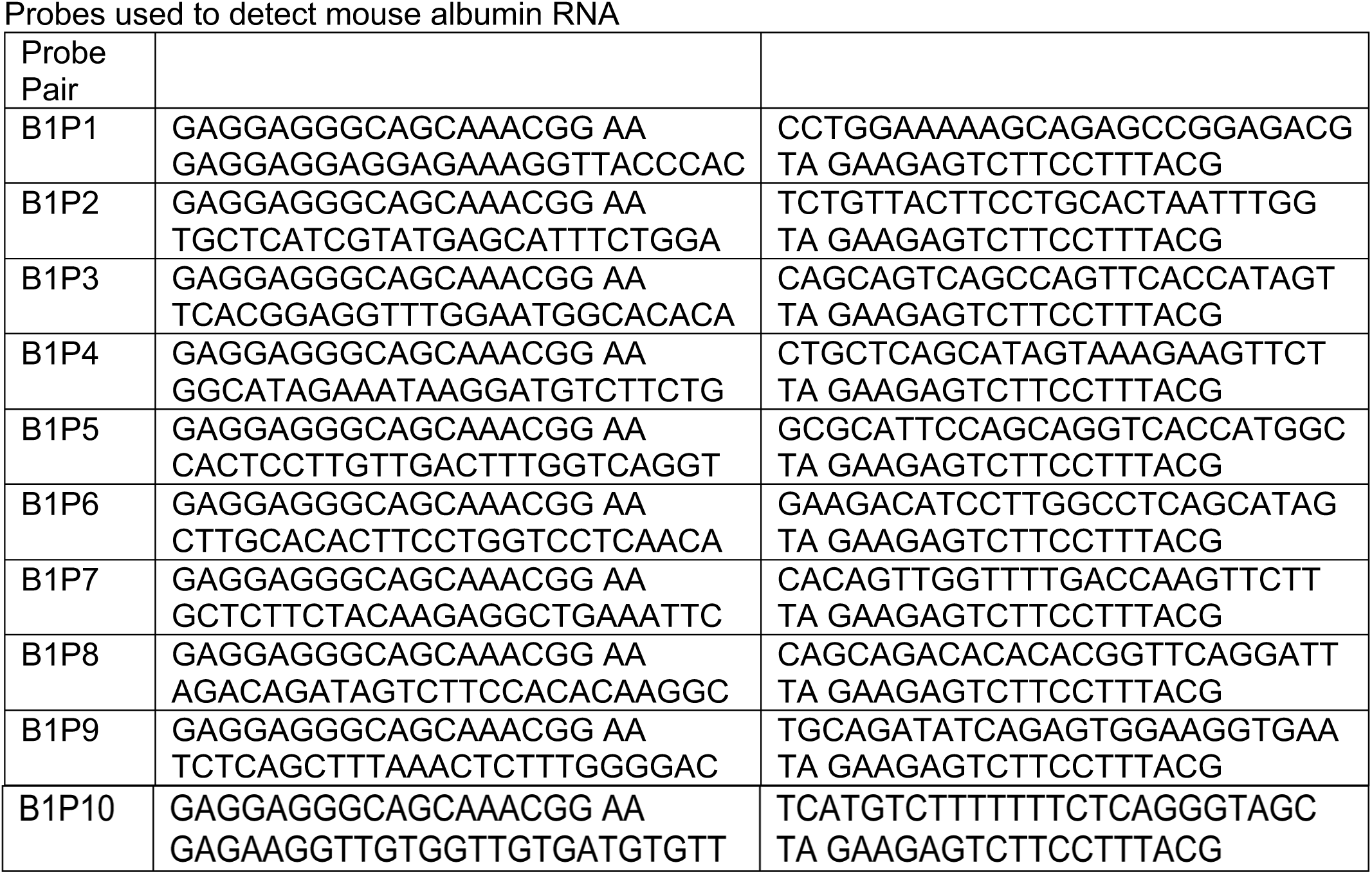

### RNA extraction and RNAseq

Total RNA was prepared using the Sigma GenElute total mammalian RNA miniprep kit with optional DNase step, according to the protocol of the manufacturer. RNA quality was assessed by Nanodrop and an Agilent RNA Screen Tape System, and 1ug was used for library preparation using RNA with Poly A selection kit (Illumina), as per the manufacturer’s instructions. Sequencing was performed on an Illumina HiSeq. RNA-sea FASTQ data were processed and mapped to the mouse reference genome (GRCm38) using CLC Genomics Workbench 20 (Qiagen). Differential gene expression was performed using the DESeq2 package in R^43^. Heatmaps were made in R using the pheatmap: pretty heatmaps package shown as the log_2_RPKM. Raw sequencing files have been deposited in Sequence Read Archives (SUB8204864, PRJNA665496).

### Luminex assays

Luminex profiling was performed on whole blood that was allowed to clot for 20 minutes and then spun down using a custom mouse IFN kit (IFN alpha, IFN beta, IL-28, Invitrogen), mouse cytokine 23-plex (Bio-Rad, M60009RDPD), and mouse chemokine 31-plex (Bio-Rad, 12009159), according to the manufacturer’s protocol. Assays were read on a Millipore MagPix machine by the Luminex Corporation. Heat maps were generated using the fold change in concentration (picograms/milliliter) of each animal compared to the average of uninfected animals and was made in GraphPad Prism. Violin plots are shown as the concentration for each animal (one point) in picograms/milliliter.

### Statistics

All statistical analysis was performed using GraphPad Prism version 8. Data are presented as mean ± SD. A one-way ANOVA was used to determine statistical significance, as described in the figure legends. Parametric tests were applied when data were distributed normally based on D’Agostino–Pearson analyses; otherwise nonparametric tests were applied. P values of <0.05 were considered statistically significant, with specific P values noted in the figure legends.

## Acknowledgements

We thank Charles Good (UPMC Children’s Hospital of Pittsburgh) and Kathryn Lemon (UPMC Cancer Center) for technical assistance, Jeffrey Bergelson (Children’s Hospital of Philadelphia) for reagents, Runjan Chetty (Brighton and Sussex University Hospitals NHS Trust) for blinded pathology analysis, and Terence Dermody (UPMC Children’s Hospital of Pittsburgh) for helpful suggestions. This project was supported by NIH R01-AI150151 (C.B.C), NIH R01-AI081759 (C.B.C.), NIH T32-AI060525 (A.I.W), NIH F31-AI149866 (A.I.W), a Burroughs Wellcome Investigators in the Pathogenesis of Infectious Disease Award (C.B.C), and the UPMC Children’s Hospital of Pittsburgh (C.B.C). This project also used the UPMC Hillman Cancer Center Tissue and Research Pathology/Pitt Biospecimen Core and Animal Facility shared resources which are supported in part by award P30CA047904.

---

**Supplemental Figure 1.**
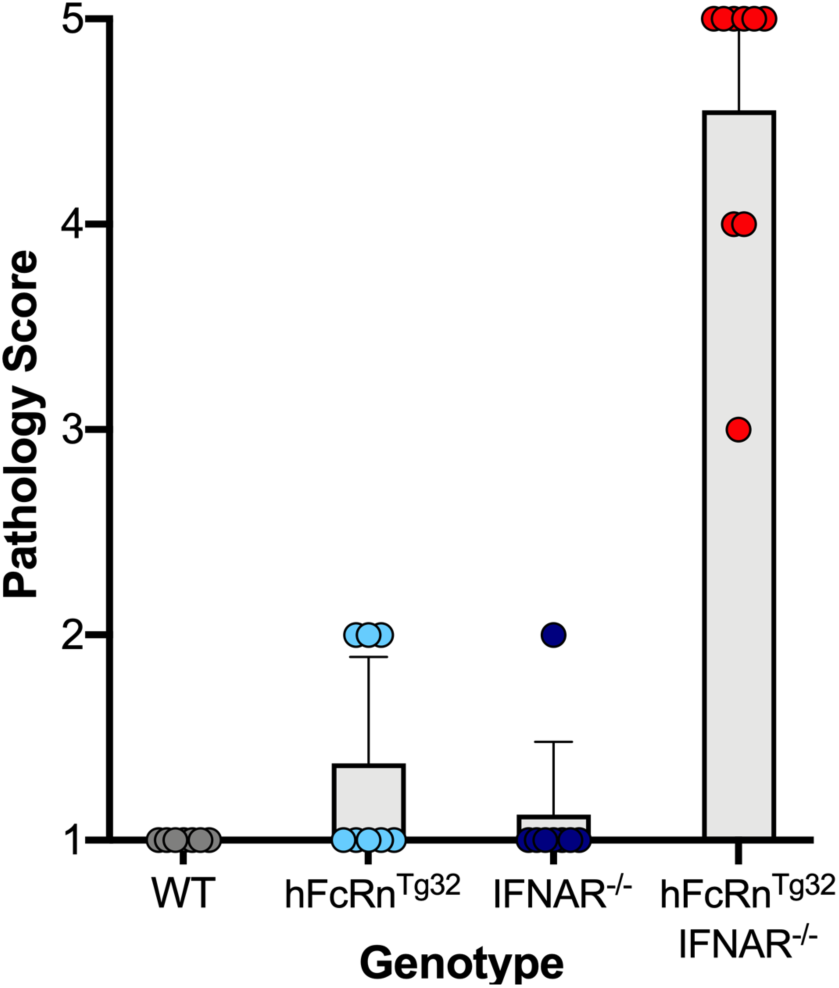
WT (grey, 7 animals), hFcRn^Tg32^ (light blue, 8 animals), IFNAR^−/−^ (dark blue, 8 animals), and hFcRn^Tg32^-IFNAR^−/−^ (red, 6 animals) adult mice were inoculated with E11 by the IP route and sacrificed 72 hours post inoculation. H&E sections were scored blinded to genotype based on severity of pathology using the following descriptors—1: retention of normal architecture and cord pattern of liver cells, 2: Immune infiltration, 3: spotty/random hepatocytolysis, 4: punctate aggregates of hepatocyte necrosis/death, and 5: confluent areas of hepatocyte necrosis and death.

**Supplemental Figure 2.**
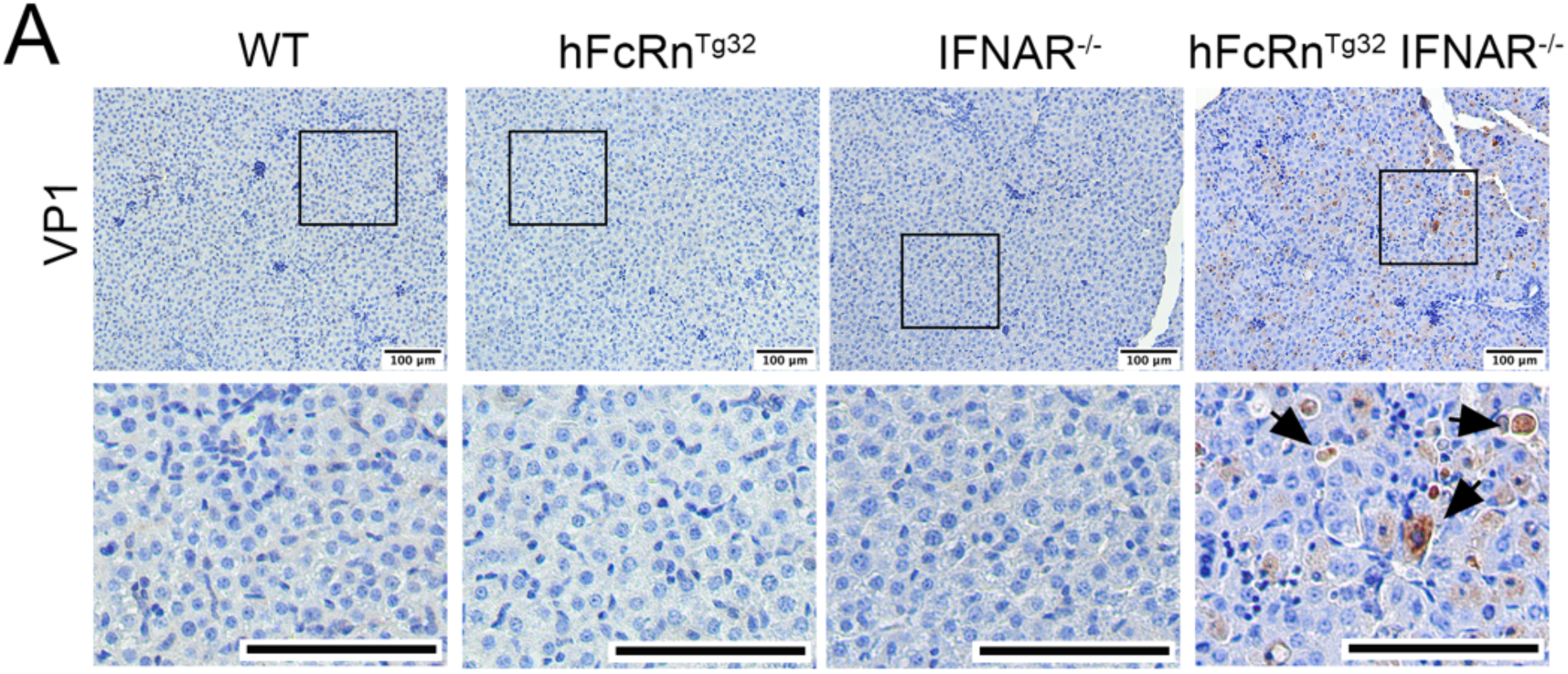
WT, hFcRn^Tg32^, IFNAR^−/−^, and hFcRn^Tg32^-IFNAR suckling mice were inoculated with E11 by the IP route and sacrificed 72 hours post-inoculation. Shown are representative images from immunohistochemistry for E11 using an antibody recognizing the VP1 capsid protein from the livers of a representative animal of each genotype.

